# VanRS and CroRS cross-talk revealed by coevolutionary modeling regulates antibiotic resistance in VanA-type vancomycin-resistant *Enterococcus faecalis*

**DOI:** 10.1101/2022.05.05.490712

**Authors:** Xian-Li Jiang, Uyen Thy Nguyen, Cheyenne Ziegler, Deepshikha Goel, Kelli Palmer, Faruck Morcos

## Abstract

*Enterococcus faecalis* is an opportunistic pathogen that can cause bacteremia and endocarditis. Previous studies have shown that concurrent treatment with cephalosporin and vancomycin antibiotics exhibit synergy in vancomycin-resistant *E. faecalis* to render the bacterium susceptible to antibiotic treatment whereas treatment with each antibiotic separately was not successful. Proteins responsible for mediating vancomycin and cephalosporin resistance are classified as two-component systems (TCS). TCS consist of a histidine kinase that phosphorylates a response regulator after environmental activation. These signaling networks have been shown to exhibit cross-talk interactions, and through direct coupling analysis, we identify encoded specificity between vancomycin resistance TCS, which are horizontally acquired, and cephalosporin resistance TCS, which are endogenous to *E. faecalis*. To verify cross-talk between these pathways is responsible for vancomycin and cephalosporin synergy, we use RNA-Seq to identify differentially expressed genes in VanA- and VanB-type vancomycin resistant enterococci after treatment with the cephalosporin antibiotic, ceftriaxone, and also with vancomycin. We find that cross-talk between VanS_A_ and CroR in strain HIP11704 may be responsible for synergy, demonstrating that horizontally acquired TCS can have large impacts on pre-existing signaling networks. The presence of encoded specificity between exogenous TCS and endogenous TCS show that the systems co-evolve, and cross-talk between these systems may be exploited to engineer genetic elements that disrupt antibiotic resistance TCS pathways.

**Author Summary:** Bacteria may transmit genetic elements to other bacteria through the process known as horizontal gene transfer. In some enterococci, vancomycin resistance genes are acquired this way. Proteins encoded within the bacterial genome can interact with proteins acquired through horizontal gene transfer. The interaction that occurs between proteins VanS_A_ and VanR_A_ is known to mediate vancomycin antibiotic resistance in VanA-type vancomycin resistant enterococci (VRE), and the interaction between proteins CroS and CroR is an important pathway in cephalosporin antibiotic resistance. We show that the VanS_A_, which is obtained through horizontal gene transfer, inhibits CroR under treatment with antibiotics vancomycin and ceftriaxone. This interaction is responsible for the observed synergy between vancomycin and ceftriaxone in VanA-type VREs. These findings demonstrate how horizontally acquired genes may produce proteins that interrupt known protein interactions, including antibiotic resistance signaling pathways in bacteria. Furthermore, the specific mechanism found for VanA-type VREs provides a basis for engineering of horizontally acquired proteins that disrupt antibiotic resistance pathways.

## Introduction

Two-component systems (TCS) are key regulatory systems for signal transduction of antibiotic-induced stress in bacteria. This pathway comprises of an environmental signal that activates a histidine kinase (HK), followed by phosphorylation of a cognate response regulator (RR) (1,2). HKs exist as integral membrane proteins and auto-phosphorylate themselves, transferring a phosphate group from ATP to a histidine residue. The phosphate group is then transferred from the histidine residue on the HK to an aspartic acid residue of the RR upon environmental stimuli. The phosphorylated RR then acts as a switch that results in diverse output responses, such as DNA-binding, RNA-binding, protein-binding, and enzymatic activities (3). While TCS interactions primarily occur between a cognate HK and RR, nonorthogonal interactions with other TCS proteins also occur (4). These non-cognate interactions impact antibiotic-induced TCS pathways and networks in bacteria, including pathogenic species and strains. In this study, we consider the interactions between horizontally acquired vancomycin-resistance TCS proteins and intrinsic cephalosporin-resistance TCS proteins in *Enterococcus faecalis* as a potential mechanism for ceftriaxone and vancomycin synergism.

*E. faecalis* is a Gram-positive bacterium that natively colonizes the gastrointestinal tracts of humans and other animals (39). It is also an opportunistic pathogen causing bacteremia and endocarditis. Infections caused by *E. faecalis* can have limited treatment options due to antibiotic resistance (4,34,35). Vancomycin is commonly used to treat Gram-positive bacterial infections and targets cell wall biosynthesis by interacting with D-alanyl-D-alanine (D-Ala-D-Ala) termini of peptidoglycan precursors. This interaction blocks the availability of D-Ala-D-Ala to transpeptidase enzymes, thereby blocking the formation of new peptidoglycan and ultimately disrupting the bacterial cell wall (5). Some vancomycin resistance genes (*van*) can be horizontally acquired by enterococci, conferring vancomycin resistance. These genes code for the synthesis of peptidoglycan precursors that terminate in D-Ala-D-lactate, for which vancomycin has low affinity. The *van* genes are tightly regulated by the VanRS TCS. This TCS exists in two transferable variants, VanA- and VanB-type in enterococci (6). Both VanA- and VanB-type enterococci confer D-Ala-D-Lactate termini, but they differ in the specific environmental signals that activate the VanRS response. The VanR_A_S_A_ TCS turns on vancomycin resistance genes in response to unknown factors associated with vancomycin-induced stress, while the VanR_B_S_B_ TCS turns on vancomycin resistance genes in response to direct vancomycin binding (6). Previously, deletion of *vanS* led to constitutive expression of *van* genes, indicating that VanS acts as a negative regulator to VanR in the absence of vancomycin. Thus, in the absence of vancomycin, VanS acts as a phosphatase, removing the phosphate group transferred to VanR by acetyl phosphate (5,6). In the presence of vancomycin, VanS then acts as a kinase, phosphorylating VanR and inducing vancomycin resistance.

VanA- and VanB-type *vanRS* are acquired by horizontal gene transfer in enterococci, which already encode endogenous and strain-variable TCS, setting the stage for potential cross-talk interactions between non-cognate TCS partners. While very little is known about how horizontal acquisition of TCS impacts existing signaling networks in *E. faecalis* or other pathogens, many examples of cross-talk between intrinsic TCS during antibiotic treatment exist (10,42). One endogenous *E. faecalis* TCS, CroRS, is particularly important for cephalosporin resistance in enterococci and exhibits cross-talk with another endogenous TCS, CisRS. The phosphorylation of CroR by CisS has been implicated in ceftriaxone antibiotic resistance phenotypes in *E. faecalis* (10,41). Interestingly, while CroRS is core to the *E. faecalis* species, CisRS is present in only a subset of strains. This subset includes a VanA-type vancomycin-resistant enterococci (VRE), HIP11704 (10,36). The intersection of intrinsic CroRS, CisRS, and extrinsic VanR_A_S_A_ in HIP11704 is of interest because synergism has been found between ceftriaxone and vancomycin in VREs (33). VanB-type *E. faecalis* V583 mediates vancomycin resistance through VanR_B_S_B_ and ceftriaxone resistance through VanY_B_ inhibition of CroRS. While VanY_B_ is responsible for the synergism between vancomycin and ceftriaxone treatment in VanB-type VREs (33), the mechanism for synergism has been unknown for VanA-type VREs. It is possible that cross-talk interactions between TCS in *E. faecalis* induce clinically relevant antibiotic resistance phenotypes, and the gain or loss of TCS proteins has significant impact on these phenotypes. By comparing VanA-type HIP11704 and VanB-type V583 response to ceftriaxone and vancomycin treatment, we may be able to uncover important TCS cross-talk interactions and their variation between strains.

Coevolutionary analysis of TCS proteins can detect potential cross-talk interactions (11,12,14). Preference for binding the right TCS partner(s) for subsequent signal transfer is driven by the recognition of a group of coevolving residues at the binding interface (8,9). When different HKs share common key coevolving residues, cross-talk is likely to happen between non-cognate pairs, such as CisS-CroR (10). Coevolution stress between the functional HK and RR domains maintains the binding preference and signal transfer in HK and RR pairs (11,12). Therefore, we applied a well-established coevolutionary computational model, direct coupling analysis (DCA) (13) to the TCS in three model *E. faecalis* strains (Table 1), one a VanA-type vancomycin-resistant *E. faecalis* HIP11704 (36), one a VanB-type *E. faecalis* V583 (37,38,40), and one a vancomycin-susceptible *E. faecalis* OG1RF (39). We use coevolution of amino acids between HK and RR sequences to investigate the impact of horizontal acquisition of TCS on existing TCS networks in these strains. DCA and its derived model are able to efficiently and accurately predict protein structures (13,14), dynamics (15) and interactions (16,17). DCA provides a global inference framework to model a joint probability distribution along the protein sequence, which is the concatenated sequence for HK and RR cognate partners. It utilizes a Boltzmann distribution to describe the probability of a naturally occurring sequence for TCS partners (Equation 1). A modified version of DCA was used to infer the two types of parameters of this distribution, pairwise coupling strength and local field. The parameters carry the information of coevolutionary coupling strength of two residue positions along the alignment sequence in the context of different amino acid settings as well as the contribution from each single position. These parameters have been used successfully to infer important connectivity between biomolecules (16,18), predict binding specificities in TCS (7,12,43) and predict functionality of proteins with rewired functional domains (19).

**Table 1.** Strains used in this study.

| Strain | Description |
| --- | --- |
| *E. faecalis* HIP11704 | VanA-type VRE |
| *E. faecalis* V583 | VanB-type VRE |
| *E. faecalis* OG1RF | Vancomycin-sensitive<br>*Enterococcus* |

In this study, a binding specificity score, ΔĤ(S), for any HK and RR pair was calculated using coevolutionary couplings at the binding interface, as found in the global Boltzmann distribution. We generated a binding specificity landscape for the TCS in *E. faecalis* to infer potential cross-talks between non-cognate pairs. During evolutionary analysis, VanR_A_S_A_ was found to possibly interact with the known CroRS-CisRS network. This interaction could be disruptive to ceftriaxone treatment response pathways. This finding was further explored via transcriptomic and mutational experiments in response to vancomycin and ceftriaxone, which are regulated by VanRS and CroRS respectively.

## Results

### Coevolutionary model predicts potential VanS_A_-CroR cross-talk in *E. faecalis* HIP11704

The coevolution-related parameters, coupling strength and local fields, from the modeled distribution of cognate and random HK-RR were inferred by a DCA-based methodology (Figure 1A). Then, we computed a binding specificity score ΔĤ(S), which is a specific version of the coevolutionary energy function of the global distribution model, for TCS pairs in the VanA-type VRE *E. faecalis* HIP11704 (Figure 1B), the VanB-type VRE *E. faecalis* V583 (Figure S1), and the vancomycin-susceptible model strain *E. faecalis* OG1RF (Figure S1). A heatmap of ΔĤ(S) was generated for all possible combinations of HKs containing HisKA domain and RRs containing Response_reg domain for each of the 3 strains, to evaluate the strength of binding specificities between different HKs and RRs. Cognate pairs exhibit much stronger binding specificity than non-cognate pairs in all 3 strains with few exceptions (Figure 1B, Figure S1), indicating the good performance of ΔĤ(S) at predicting binding partners in the TCS family. HIP11704 and OG1RF share most HK and RR proteins, with HKRR08 and VanRS_A_ only found in HIP11704, and RR01 and RR15 in OG1RF. Most importantly, CisRS and CroRS homologs are present in both HIP11704 and OG1RF, and OG1RF does not have *van* genes. There is no difference for ΔĤ(S) scores between shared nodes in HIP11704 and OG1RF, as all the differential edges are related with the non-shared node (Figure 2A). However the introduction of VanRS_A_ creates new strong connections, especially with CroRS and CisRS. In the differential network comparison between HIP11704 and V583, there are more altered connections. Signaling networks between VanRS and other TCS proteins are different in HIP11704 and V583 (Figure 2B), consistent with the fact that these two strains confer distinct VanRS TCS (6). VanS_A_ in HIP11704 shows enhanced specificity with other non-cognate RRs, while VanR_A_ in HIP11704 has higher specificity towards its cognate partner, VanS_A_ (Figure 1B, S1 and 2B). CisRS is not found in V583 and causes strong rewired connections with VanR_A_S_A_ in HIP11704. These differential networks show that new TCS or different TCS orthologs introduce novel and strong connections in existing TCS signaling networks. These new signaling pathways may lead to differential phenotypes.

**Figure 1.**
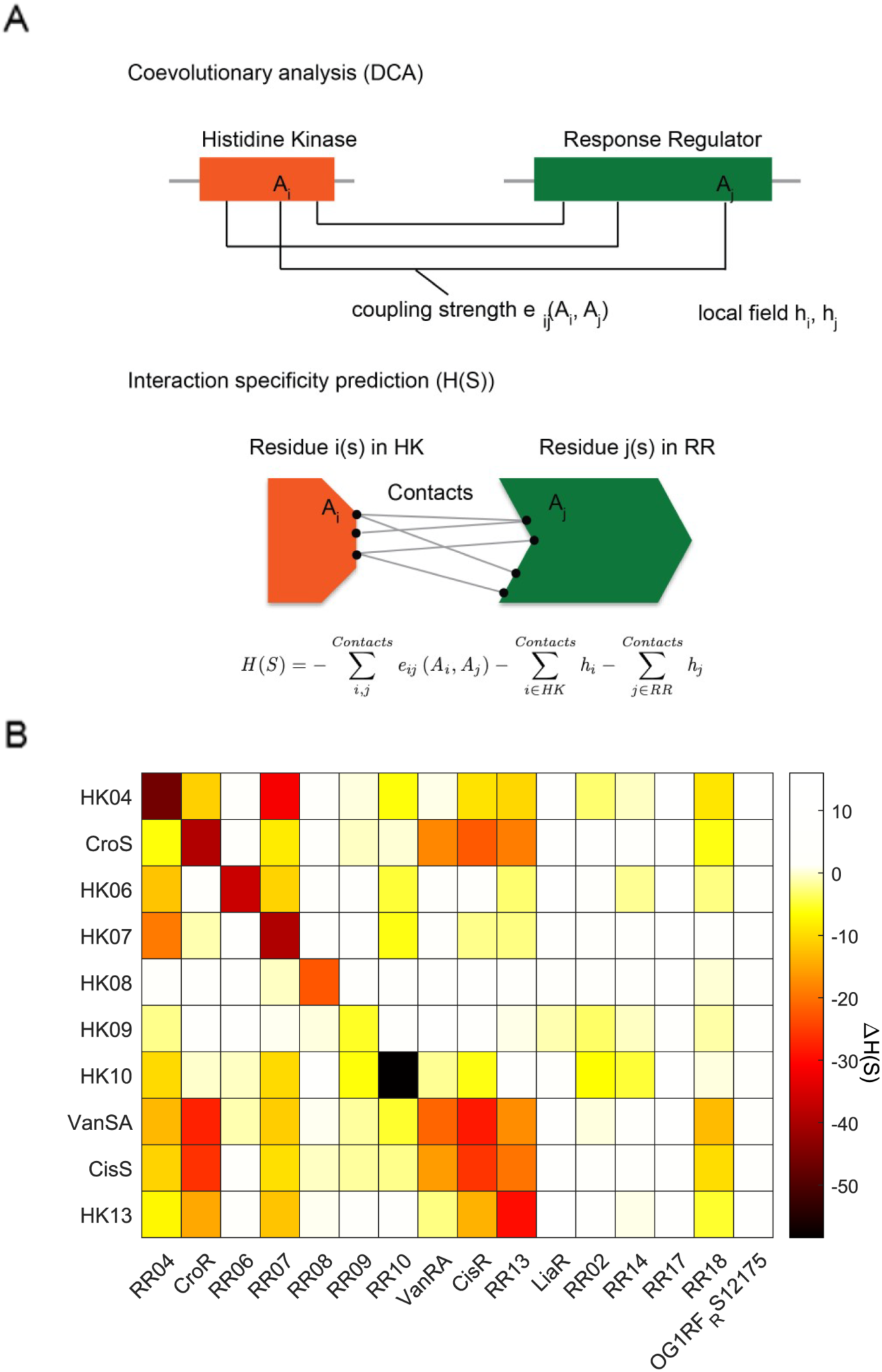
Coevolutionary model predicts binding preference among HKs and RRs in *E. faecalis*. A. Binding specificity score calculated from coevolutionary model based. B. Binding specificity landscape of 10HKs and 16 RRs in *E. faecalis* HIP11704. A lower ΔĤ(S) score with a darker color indicates stronger binding preference.

**Figure 2.**
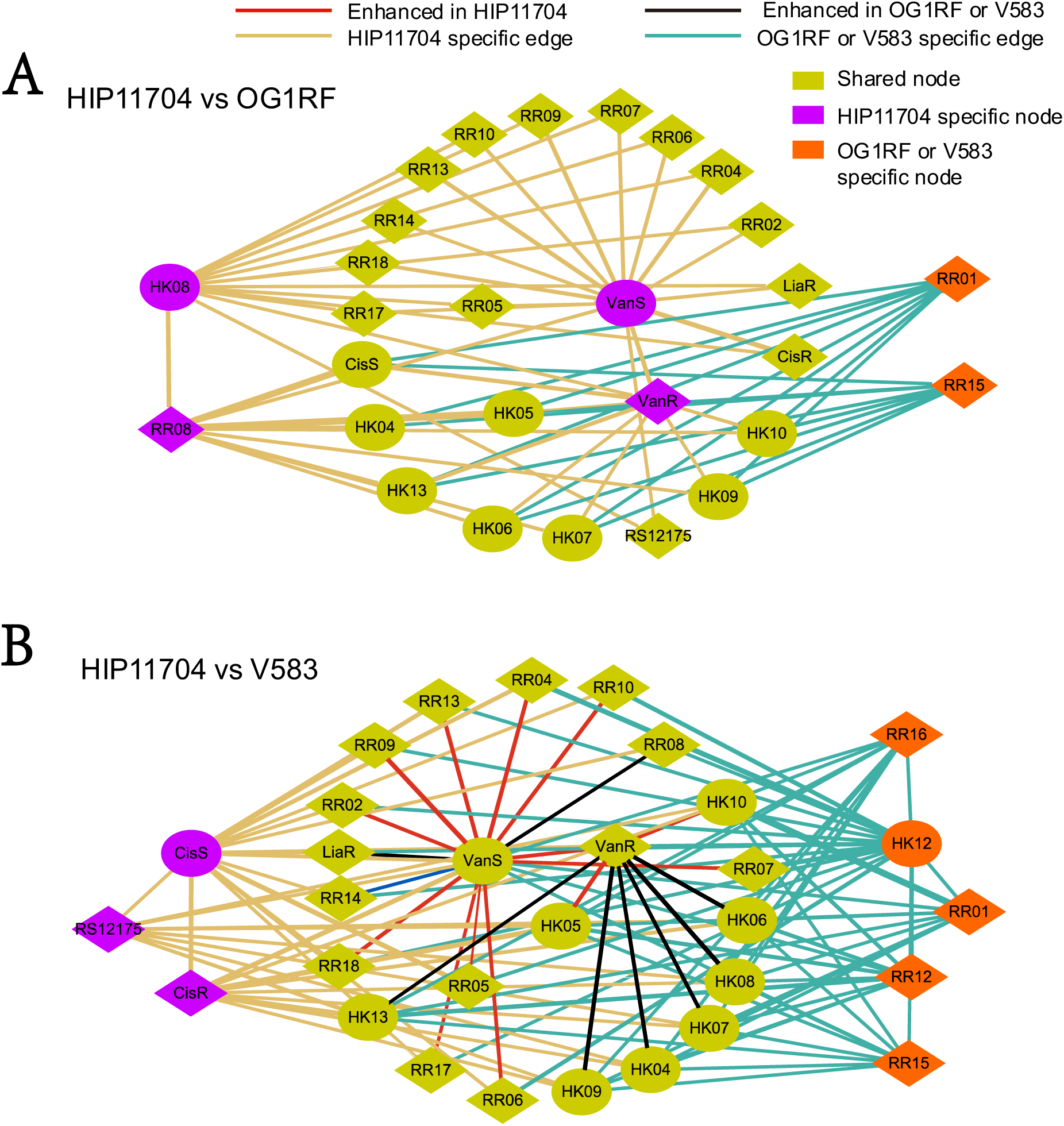
Differential TCS network between HIP11704 with OG1RF (A) or V583 (B). Each node represents a HK or a RR, with purple color specific for proteins in HIP11704 and orange color specific for V583. Each edge indicates that there is a difference of HK-RR binding specificity scores, ΔĤ(S), between two strains with the thicker ones representing larger difference.

Certain cross-talk signals from the heatmap (Figure 1B) have been previously identified. HK04 has a comparable binding specificity with RR07, and their homologous proteins, PhoR and YycF in *B. subtilis*, exhibit crosstalk (28). Another cross-talk interaction is found between CroRS (HKRR05) and CisRS in HIP11704 (Figure 1B) and OG1RF (Figure S1A), which was also confirmed in recent studies (10). For the potential cross-talk between CroRS and HKRR13, which had been excluded by a previous study (10), CroS-CisR shows stronger specificity strength than CroS-RR13, and CisS-CroR is stronger than HK13-CroR (Figure 1B, Figure S1A). Additionally, VanS_A_-CroR has a stronger score than CisS-CroR and its cognate partner VanS_A_- VanR_A_, indicating potential interference to the verified cross-talk between CroRS-CisRS. Such inference is absent in OG1RF, which lacks VanS_A_ and VanR_A_, and in V583, which carries VanS_B_ and VanR_B_ instead.

The deletion of CroS led to the constitutive phosphorylation of CroR in strains containing CisRS (10). However, this was only confirmed in vancomycin-susceptible strains and a VanB-type vancomycin-resistant strain. The rationale behind putative binding between CisS and CroR is supported by the fact that CisS and CroS share 6 residues out of 9 total residues that are key to binding specificity. To explore potential competition between CisS and VanS_A_ for CroR, we focused on the same 9 key specificity residues on these three HKs (7-9). A comparison of those 9 residues between VanS_A_ and CroS also shows 6 of them are shared by VanS_A_ and CroS, while the number of common key residues between CisS and CroS is 5. VanS_B_ shares none of the 5 CisS-CroS common residues (Figure S2). We propose that VanS_A_ may act as a competitor of CisS in the absence of CroS (Figure 3) when VanR_A_S_A_ and CroRS regulate resistance to vancomycin and ceftriaxone, respectively. Vancomycin has been found to trigger the CroRS expression via an unknown mechanism, which may be directly acting on CroS (40). Additionally, among 12 published specificity residues (8,9) in RRs, VanR_A_ and CisR share 9 residues with CroR, while RR13 shares 8 residues and VanR_B_ shares 5. Cumulatively, these results suggest that cross-talk between CroRS and CisRS may be affected by the presence of VanS_A_ and VanR_A_. Thereby antibiotic resistance behavior, especially the hyper-resistance in the *croS* mutant strain, might be different in HIP11704. Further evidence is needed to characterize the role of VanS_A_ to CroR.

**Figure 3.**
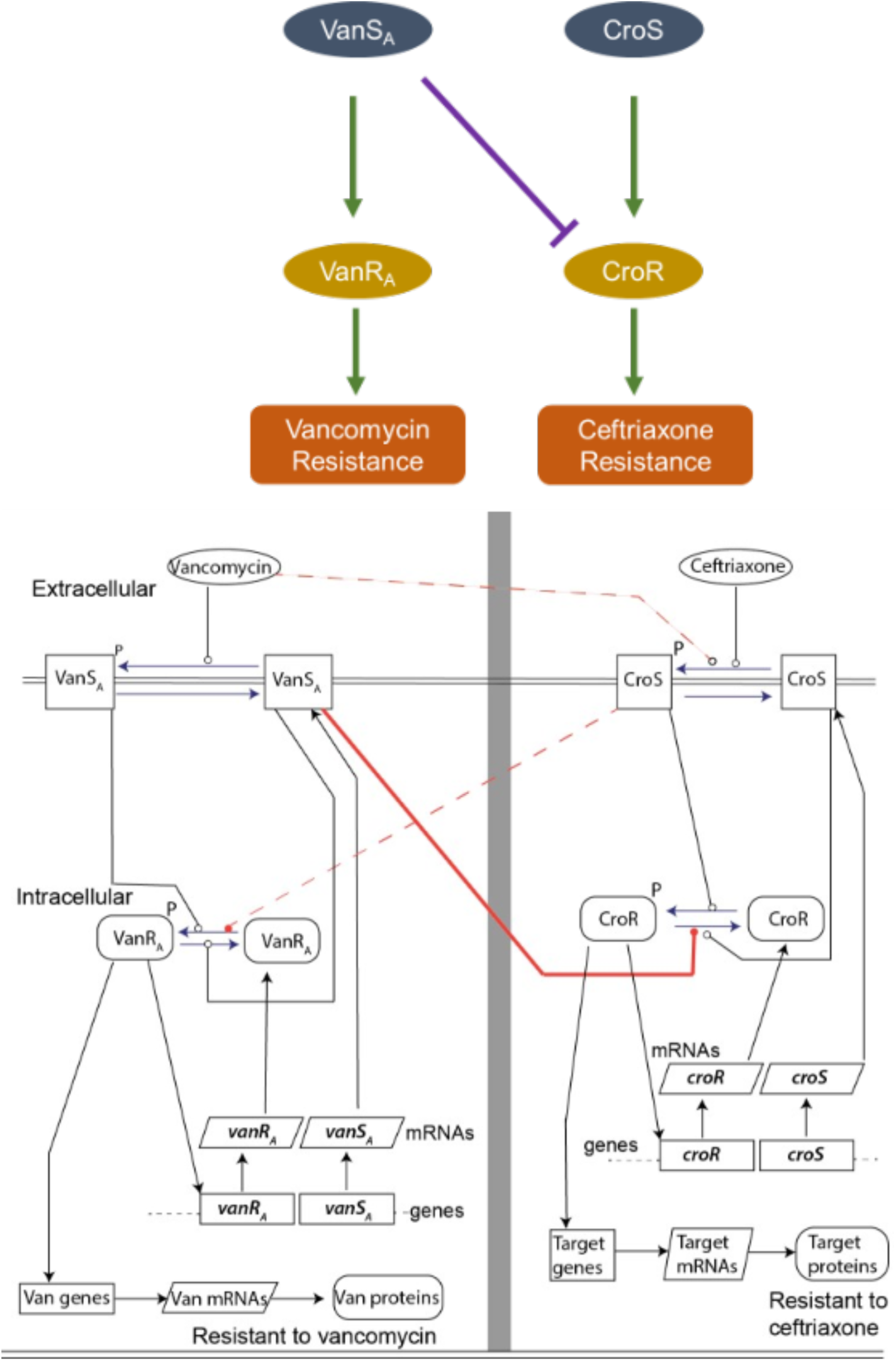
Possible cross-talk between VanR_A_S_A_ TCS and CroRS TCS. In HIP11704, VanR_A_S_A_ and CroRS regulate the resistance to vancomycin and ceftriaxone, respectively. Red solid line indicates the main cross-talk path explored in this study. Dashed lines represent two other possible cross-talk paths, where the CroS-VanR_A_ together with VanS_A_-CroR constitute a reciprocal VanR_A_S_A_-CroRS cross-talk network.

### Strain-specific and cell-wall targeting antibiotic-specific alterations of TCS detected through RNA-Seq

TCS are important to the response of bacteria to external stimuli, such as antibiotic-induced stress. We have shown that the binding specificity landscapes of TCS are different between HIP11704 and V583 (Figure 1B). To explore gene activation patterns in TCS, we performed RNA sequencing to examine differential gene expression in V583 and HIP11704 in response to vancomycin and ceftriaxone exposures at MIC. Our rationale was that different cross-talk interactions between TCS should lead to different transcriptomic responses in these two strains. This landscape is also captured when considering co-expression of HKs and RRs over all treatment groups as measured by RNA-Seq (Figure S3). When comparing binding specificity score to log fold change correlations, we find that noncognate HK-RRs with very strong specificity scores all exhibit co-expression (Figure S4). Agreement to the model weakens as scores predict less specificity, which is expected since predicted binding specificity only considers aligned residues within the interface and does not account for the unaligned portion of each TCS protein.

HIP11704 and V583 share 15 homologous HKs and 14 homologous RRs. Transcriptomic changes in these TCS HK and RR genes were also different across strains and treatments (Figure 4). Unlike V583, HIP11704 contains CisRS, which showed a log fold increase during both ceftriaxone and vancomycin treatment (Figure S6). HIP11704 TCS were more responsive than V583 and more responsive to vancomycin than ceftriaxone (Figure S5). Notably, *ef*1820 (HK15) was inversely regulated in HIP11704 and V583 under vancomycin treatment. *ef*1820, known as *FsrC*, is a part of fsr quorum-sensing locus that regulates biofilm development by producing gelatinase and a serine protease (29,30). However, no homologous RR15, cognate partner for HK15, was found in HIP11704. Some TCS genes exhibited differential expression across strains and antibiotic treatments. For example, HKRR06 were upregulated in both V583 and HIP11704 for vancomycin, indicating involvement of HKRR06 in vancomycin-induced response. HKRR10 were down-regulated by vancomycin in HIP11704 and remained unaltered in V583. The binding specificity profiles for HKRR10 were different in V583 and HIP11704 due to the introduction of HKRR12 in V583 (Figure S1B and Figure 1B). HKRR09 also exhibited V583 specific down-regulation in response to vancomycin. The TCSs binding specificity landscape explicitly shows changes in TCS binding specificity profile after introduction of new or orthologous TCS proteins. These changes in specificity may provide an explanation for distinct expression behaviors across strains.

**Figure 4.**
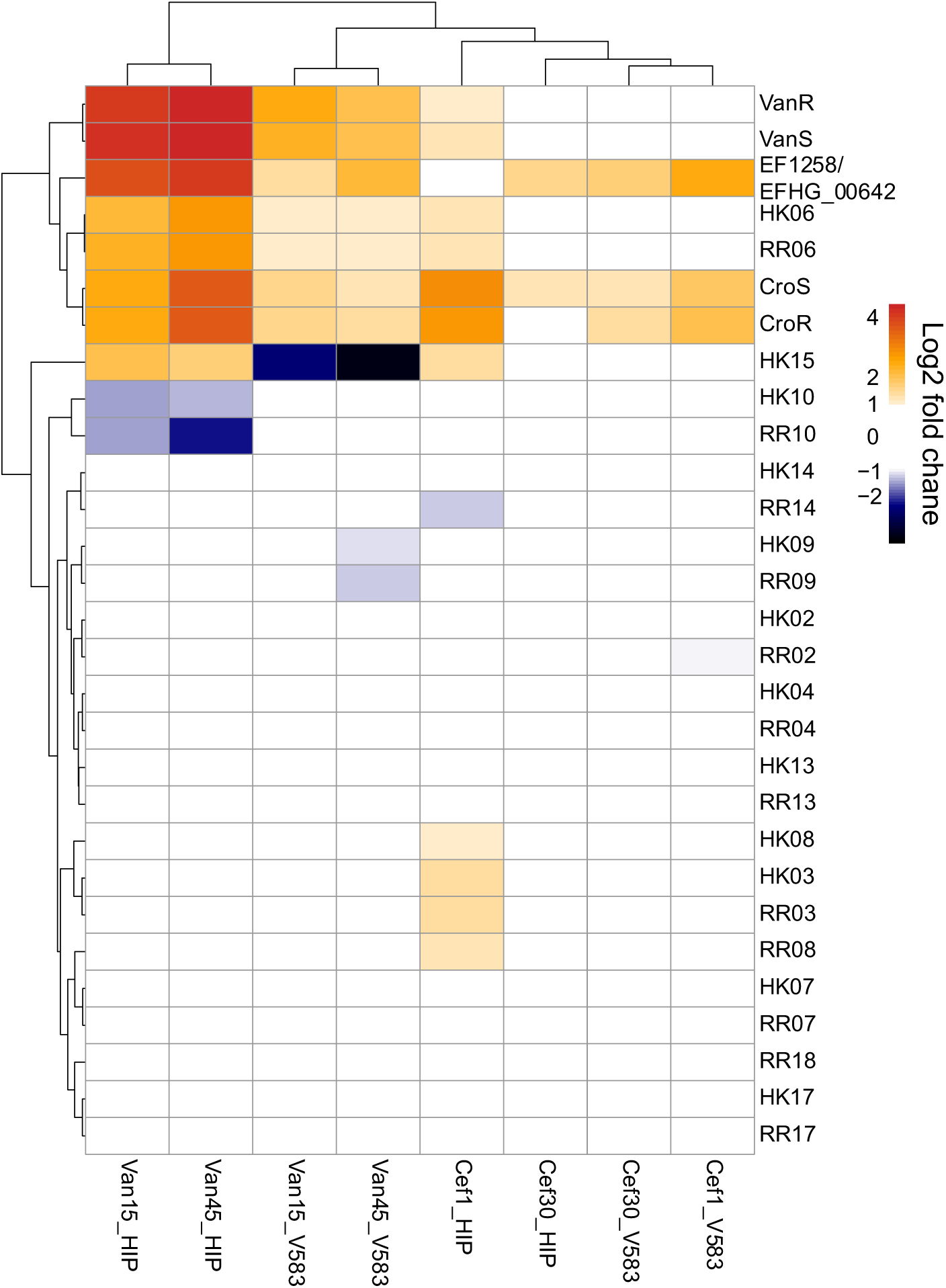
Fold change of TCS genes in HIP11704 and V583 under 4 conditions. Log 2 of fold change levels between -1 and 1 are shown in white color, while more red color indicates up-regulation and blue indicates down-regulation. No FDR information was included in this heatmap.

Expression values of *vanRS* and *croRS*, which are responsible for the resistance to vancomycin and ceftriaxone respectively, and their target genes are shown in Figure 5 and Figure S7. Both vancomycin and ceftriaxone induced the upregulation of *croRS*. Meanwhile, the known downstream, *salB* (*ef*0394) (31) and *glnQ* (*ef*1120) (32) in V583 remained unchanged, and for HIP11704, the *glnQ* ortholog remained unaltered and *salB* was down regulated. *cisRS* and its immediate downstream genes are highly upregulated in response to vancomycin but not in response to ceftriaxone (Figure S8). Genes in the *van* operon of HIP11704 and V583 were both highly upregulated when exposed to vancomycin, as expected. As shown in Figure 5, the VanR_A_S_A_ was induced more than VanR_B_S_B_, while the induction level of Van_B_Y_B_WX was much higher than the Van_A_X_A_Y_A_. This indicates that VanB type vancomycin-triggered response is more efficient than VanA type. Ceftriaxone induced a mild upregulation of Van_A_X_A_Y_A_ in the context of unchanged level of VanR_A_S_A_. On the other hand, the VanB type operon (Van_B_Y_B_WX genes) did not respond to ceftriaxone. The mechanism for differential expression of VanA and VanB operons under ceftriaxone may be indicated by the comparable binding signal between CroS-VanR_A_ (Figure 1B) and the weak signal for CroS-VanR_B_ (Figure S1B).

**Figure 5.**
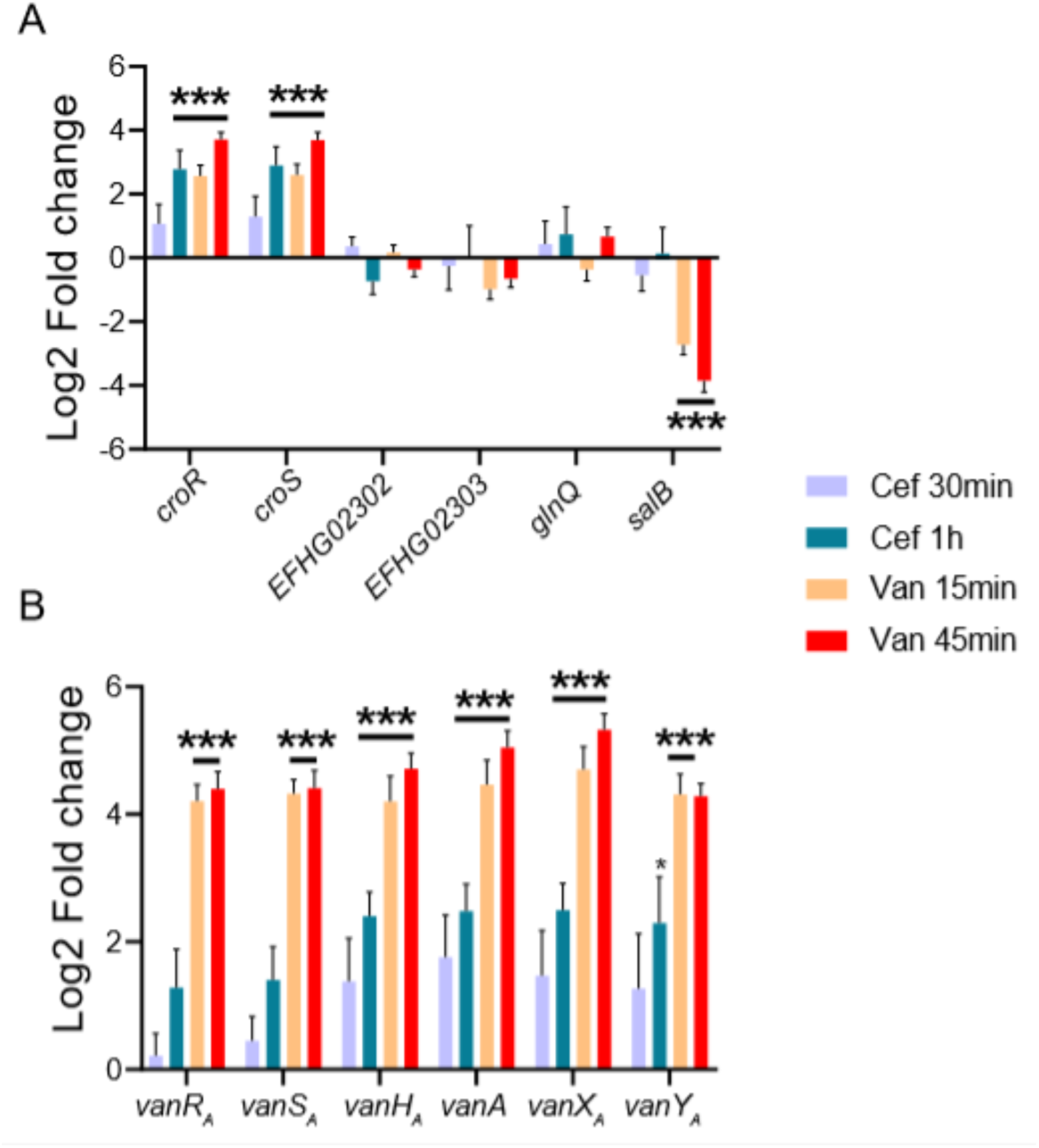
Transcriptomic changes of *vanRS* and *croRS* and their (potential) target genes in HIP11704. A. CroRS and their potential targets. *EFHG_02302* and *EFHG_02303* are encoded directly downstream of *croR. glnQ* and *salB* were previously shown to be regulated by CroRS but are not responsible for ceftriaxone resistance (32). B. Expression levels of genes in *van* operon. Genes are arranged in the order of transcription direction. One star indicates p<0.05 for test group, two stars indicates adjusted p<0.05 for test group, and three stars indicates adjusted p <0.05 and passes Bonferri test for the test group.

There is synergism between vancomycin and ceftriaxone both in V583 and HIP11704, with VanY_B_ mediating synergism in V583 (33). Our results show, in V583, vancomycin-induced VanY_B_ is upregulated, CroRS is upregulated, but transcription product of CroR, *salB* is downregulated (Figure S7). Thus, VanY_B_ inhibits the CroRS pathway, and in the presence of both vancomycin and ceftriaxone, confers susceptibility through this mechanism. With the presence of high transcription levels of *vanY*_*A*_, HIP11704 still inhibits resistance to ceftriaxone (Figure 5). This indicates that synergism in HIP11704 is mediated in a VanY_A_-independent manner. The distinct responses to ceftriaxone in the context of CroRS may be attributed to the different TCS interaction networks (Figure 1 and Figure 2).

### Differential transcriptomics response to two different cell-wall targeting antibiotics in Van-A and Van-B type VREs indicates notable differences in mechanism between VREs

We analyzed global gene expression patterns in the antibiotic-treated cells to discern differences between different VRE types. Genetic expression changed more in HIP11704 than V583 when treated with vancomycin, whereas ceftriaxone treatment triggered more changes in V583 (Figure 6). As exposure time increases, may changes in expression may be attributed to changes in growth rate (Figure S9). During ceftriaxone treatment, few genes exhibited significant changes in levels of expression for both V583 and HIP11704, with *croRS* being strongly upregulated in both strains. In contrast, both strains exhibited widespread changes in expression when treated with vancomycin. Under vancomycin treatment, HIP11704 experienced strong upregulation of *vanR*_*A*_*S*_*A*_ and V583 experienced strong upregulation of *vanB, vanR*_*B*_*S*_*B*,_ and *vanWXY*_*B*._ The lower number of differentially expressed genes (DEGs) induced by ceftriaxone indicate that *E. faecalis* is less responsive to ceftriaxone than to vancomycin at the concentrations used.

**Figure 6.**
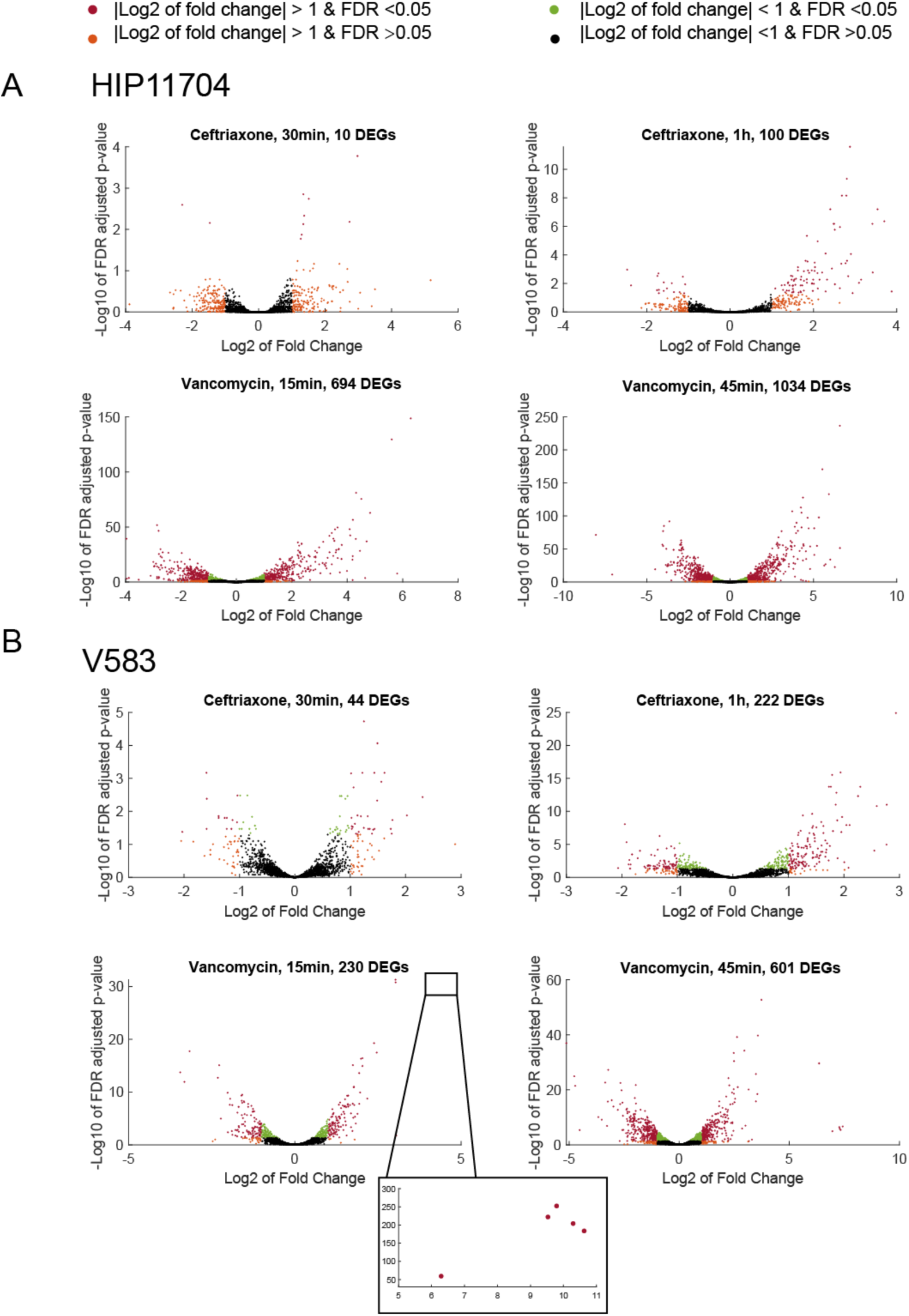
Volcano plots of transcriptomic changes in VREs under ceftriaxone or vancomycin. A. Transcriptomic changes in HIP11704. B. Transcriptomic changes in V583. For each antibiotic, there were two different treatment durations, representing short-term and long-term exposure. Differentially expressed genes (DEGs) are genes with fold change > 2 and FDR < 0.05.

To explore differences in genetic response between HIP11704 and V583, DEGs in HIP11704 were mapped to the orthologous genes in V583 (Figure 7A). Among the orthologous DEGs of HIP11704, only a fraction were regulated similarly in V583. This proportion is significantly reduced during vancomycin treatment. However, 52 of 737 DEGs in V583 possessed no homologous genes in HIP11704. Genes *ef*1058 and *ef*1057, encoding a universal stress protein and Mn2+/Fe2+ transporter, were upregulated in HIP11704 and downregulated in V583 when treated with vancomycin for 15 minutes. When the treatment increased to 45 minutes, *ef*1820, encoding HK15/ FsrC, *ef*3085, encoding an iron ABC transporter permease, and *ef*0823 were regulated inversely in HIP11704 and V583.

**Figure 7.**
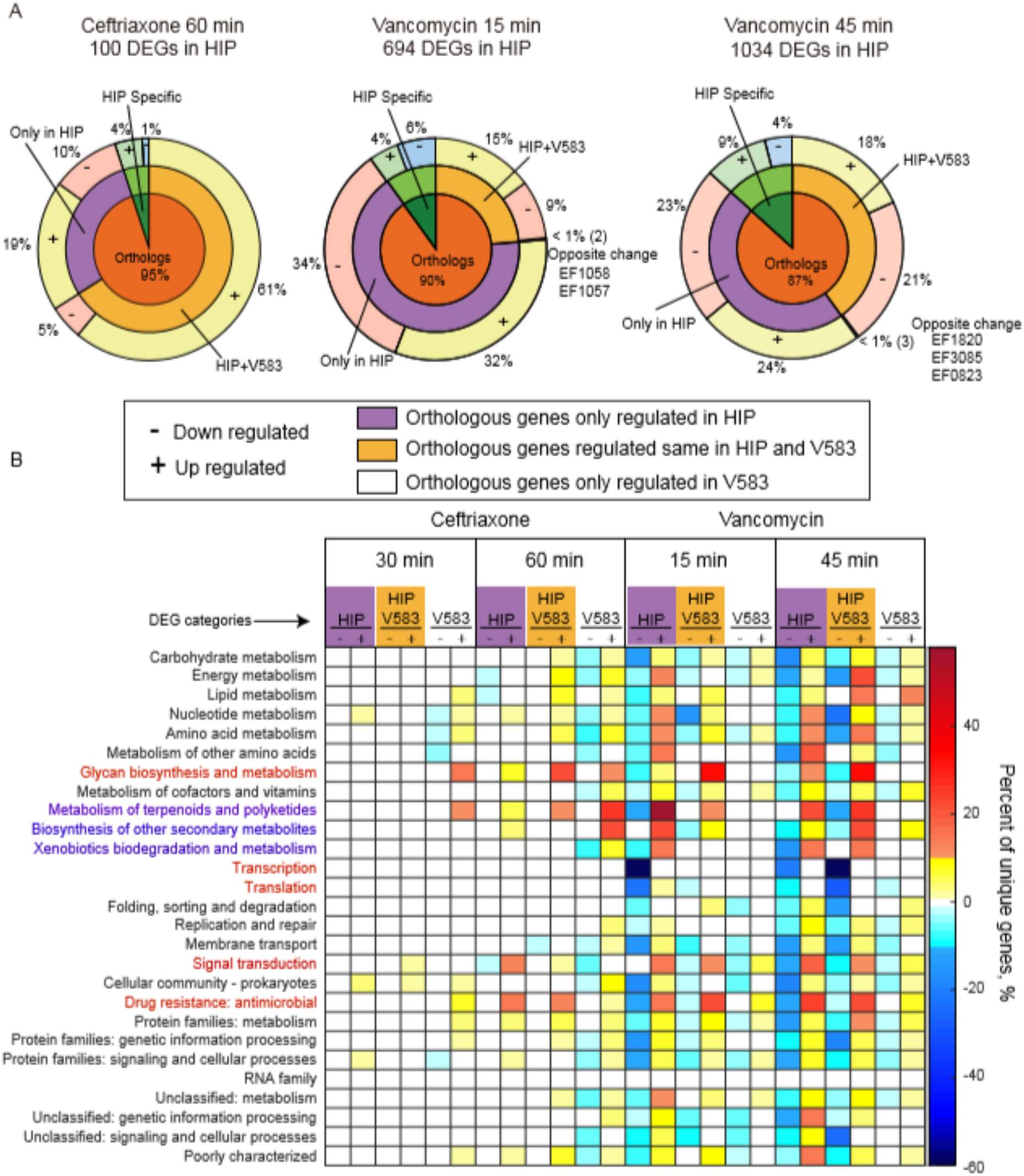
Comparison of DEGs and enriched pathways in HIP 11704 and V583 in response to vancomycin and ceftriaxone. A. Compositions of differentially expressed genes (DEGs) in HIP11704 treated with ceftriaxone or vancomycin shown in donut-pie chart. In the first layer-donut, DEGs are divided into 2 classes: Orthologous genes and genes only can be found in HIP11704. In the second layer–donut, orthologous genes are further divided into 2 or 3 classes: genes regulated only in HIP, genes regulated same in HIP and V583, gene regulated but oppositely in HIP and V583. Each block in the second layer – donut is further divided into 2 classes to make the third layer: up regulated and down regulated. The percent numbers next to the third layer donut blocks represent the proportions of the number of this DEG category to the total DEGs induced by the treatment in HIP11704. B. Enrichment of KEGG orthology (KO) for different DEGs category for four treatments. All the orthologous genes in HIP and V583 were analyzed against the KO database. There are 3 DEG categories for each treatment: only changed in HIP11704, both changed in HIP11704 and V583, only changed in V583 and then each category is divided into down regulated or up regulated. Color bar was used to indicate the percent of genes changed in each function orthology. After excluding the KO term ‘Not included in regular maps’, 27 KOs were found to be related with V583. In V583 and HIP11704, among the DEGs, about 62% and 63% genes could be annotated by KO, while the KO information of the rest genes were missing. Then for each KO, the percent of genes found in each DEG category was calculated by dividing the number of genes belonging to this KO and this DEG category by the total number of genes in this KO. Down regulated genes are shown in negative value and blue colors, up regulated genes are shown in positive and yellow-red colors.

Kyoto Encyclopedia of Genes and Genomes (KEGG) analysis was conducted for the orthologous DEGs induced by ceftriaxone and vancomycin treatment at two different time points. KO terms were classified by orthologous DEG category: only regulated in HIP11704, only regulated in V583, and regulated in both in HIP11704 and V583 (Figure 7B). Ceftriaxone treatment induced the upregulation of specific glycan biosynthesis and metabolism by 60 minutes in HIP11704 orthologs, V583 orthologs, and shared orthologs. However, when treated with vancomycin, V583 only orthologs exhibited no changes in regulation, indicating ceftriaxone and vancomycin cause different effects on glycan biosynthesis and metabolism in V583 and HIP11704. Downregulation in transcription and translation after 15 minutes of vancomycin treatment only occurs in HIP11704, and after 45 minutes, occurs in both HIP11704 and V583. Ceftriaxone treatment does not induce changes in expression of genes annotated for transcription and translation in either strain. For genes involved in drug resistance, 1 hour of ceftriaxone induces high levels of upregulation in HIP11704 only. Some genes are only down regulated in HIP11704 in response to vancomycin and other genes are more up regulated in HIP11704. When treated with vancomycin, HIP11704 orthologs exhibit high levels of both up- and down-regulation, whereas V583 orthologs show minimal changes in regulation. Therefore, HIP11704 and V583 exhibit differential expression to the same cell wall targeting antibiotics when considering the number of genes regulated and molecular pathways affected.

To create a profile of expression for HIP11704 and V583 under ceftriaxone and vancomycin treatment, DEGs were clustered using universal manifold approximation and projection (UMAP). In the profiles created, cognate TCS pairs are nearby and CisRS-CroRS cross-talk in HIP11704 is shown by CisRS and CroRS belonging to the same cluster (Figure 8A-B). Since known interactions are clustered and distances in UMAP are Euclidean, we see that during vancomycin treatment of HIP11704, *vanS*_*A*_ is co-expressed with RR10 (Figure 8B). This is supported by Figure S3B, which shows a strong negative correlation between the expression of *vanS*_*A*_ and RR10. Co-expression of *croRS* and *cisRS* is maintained under vancomycin treatment of HIP11704 with both TCS clustered separately from *vanR*_*A*_*S*_*A*._ During ceftriaxone treatment of V583, we see a clear separation between *vanY*_*B*_ and *croRS*, showing that VanY_B_ acts independently from CroRS. *croRS* and *hkrr06* are clustered, but co-expression is not supported (Figure 8C and Figure S3B). When V583 is treated with vancomycin, there is clustering of *croRS* and *vanR*_*B*_*S*_*B*_ which is corroborated by negative co-expression (Figure 8D and Figure S3B). Furthermore, v*anY*_*B*_ is activated under both ceftriaxone and vancomycin treatment but is separated into a small cluster during vancomycin treatment. Since it is responsible for directly binding vancomycin, this indicates that much of vancomycin resistance processes in V583 are decoupled from *vanY*_*B*_ itself and occur downstream. The analyses of the RNA-Seq data show that acquired TCS can directly impact expression and change the way TCS behave and are regulated. It is also apparent that these changes influence expression of downstream genes and other genes involved in cellular response, allowing important systems to either be downregulated or upregulated.

**Figure 8.**
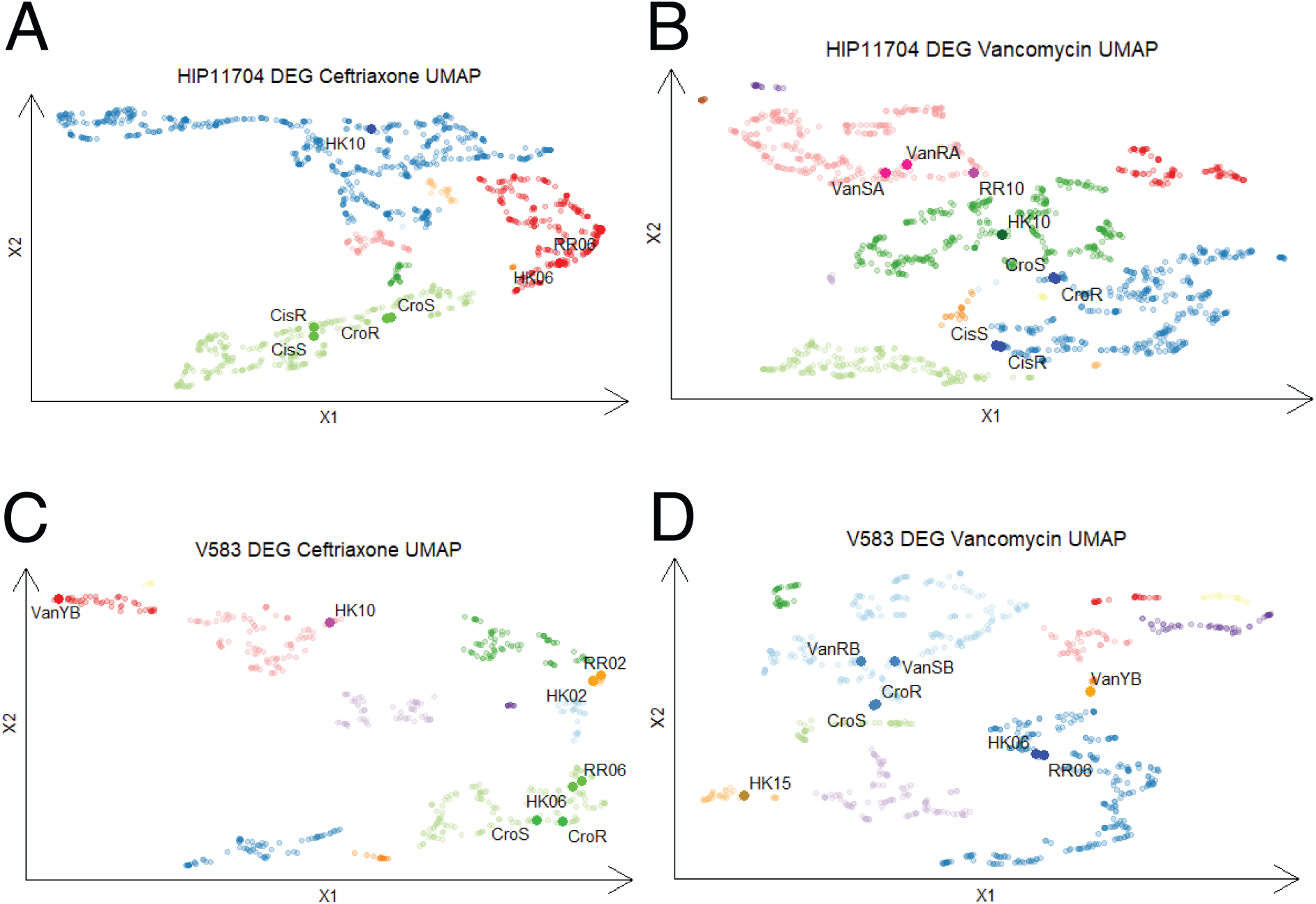
DEGs are clustered using UMAP across treatments and strains. Cognate TCS pairs are frequently nearby each other in clustering profiles, indicating that nearby proteins in each cluster are likely co-expressed, either positively or negatively. Axes define UMAP latent space. A. HIP11704 DEGs after treatment with ceftriaxone across all time points. B. HIP11704 after treatment with vancomycin across all time points. C. V583 DEGs after treatment with ceftriaxone across all time points. D. V583 DEGs after treatment with vancomycin across all time points.

## Discussion

In this study, we investigated the interaction between proteins CroRS, CisRS, and VanR_A_S_A_, which show strong reciprocal preference, or cross-talks, in binding for phosphotransfer and phosphatase activities. These three systems simultaneously exist in VanA type HIP11704 to create antibiotic-resistance synergism, while the CroRS-CisRS cross-talk is the mechanism of *croS*-deletion induced hyper-resistance in non-VanA-type systems (10). Given these predictions and observations, a potential cross-talk model based on amino acid coevolution was proposed to predict specificity between VanR_A_S_A_ and CroRS TCSs (Figure 3). The resulting model showed CroRS-CisRS cross-talk was interfered by the presence of VanR_A_S_A_ in VanA type strains. We propose that VanS_A_ acts as a phosphatase to CroR. Therefore, the CisS-dependent hyper-resistance in *croS* mutant is not observed in HIP11704. On the other hand, CroS might act as a kinase, phosphorylating VanR_A_ and conferring the upregulation of Van-A type genes, *vanH*_*A*_, *vanA, vanX*_*A*_, *vanY*_*A*_ (Figure 5). However, the VanR_A_S_A_ was not induced. This indicates that CroRS may interfere with the *van* operon in a VanR_A_S_A_-independent manner, leading to the inconclusive role of the CroS-VanR_A_ path.

VanS_A_ and CroR cross-talk may also play an important role in synergism since we propose that the role of VanY in the sensitization of enterococci to ceftriaxone induced by a subinhibitory amount of vancomycin might not account for the synergism in HIP11704. In our VanS_A_-CroR cross-talk model, vancomycin induces a much larger amount of VanS_A_, which competes with CroS and dephosphorylates more CroR. To explain the observation in *vanR* mutant under treatment of ceftriaxone plus vancomycin, we must address how the MICs were decreased when there is no more *vanS*_*A*_ produced since the transcription regulator *vanR*_*A*_ is deleted. We propose that VanS_A_ serves as a phosphatase for VanR_A_ without vancomycin but changes to kinase activity when vancomycin is present. Without a phosphoryl recipient, VanR_A_, the reversible conversion between auto-phosphorylated and dephosphorylated VanS_A_ may be biased towards dephosphorylation.

By unraveling interactions between CroRS, CisRS, and VanR_A_S_A_, we uncover mechanisms governing synergism between beta-lactam antibiotics and non-beta-lactam antibiotics in enterococci. Specifically, *E. faecalis* natively colonizes the gastrointestinal tracts of humans and other animals and are opportunistic pathogens that cause bacteremia and endocarditis (34). Infections caused by VRE have limited treatment options. Synergism between vancomycin and ceftriaxone provides new insights in treating VRE. The possible interference of VanR_A_S_A_ to the known CroRS-CisRS network is a relevant hypothesis and the presence of this interaction indicates that horizontally transferred antibiotic-resistance genes may impact endogenous signaling pathways in other pathogenic bacteria and strains. Further characterization of VanR_A_S_A_ and CroRS cross-talk not only provides a better understanding of the antibiotic resistance in Van-A type *E. faecalis*, but also may bring new insight on managing multidrug resistant infections.

## Materials and Methods

### Dataset for HK and RR functional domains

HMM profiles of HisKA domain (Pfam: PF00512) and Response_reg domain (Pfam: PF00072) were obtained from PFAM (20) and were used to search against the Uniprot database (21) using HMMER (22) command *hmmsearch* to get the multiple sequence alignments (MSAs) datasets of HK (with HisKA domain) or RR proteins. Only the sequences with E-value ≤ 10 were included in the output FASTA datasets including ∼ 270,000 HK sequences and ∼520,000 RR sequences. Each HK MSA had a length of 64 residues and each RR MSA had a length of 112 residues. The initial datasets were processed to remove obsolete sequences and hybrid protein sequences containing both HisKA and Response_reg domains. Since bacterial cognate HK and RR pairs are typically encoded adjacent in the genome and controlled by the same operon, the sequences of HK were concatenated with sequences of RR when their gene locus numbers were 1 apart, while the HKs without any genetically adjacent RR were excluded from the dataset. The resulting cognate HK-RR sequence was denoted as S=(A_1_, A_2_,…, A_n_), where n is the length of the concatenated MSA and equals to 176, A represents the aligned value, selected from 20 amino acids or 1 alignment gap, at certain position. The final dataset, hereinafter referred as HK-RR dataset, contains 74,451 sequences and was used as the input dataset for coevolutionary analysis for TCS partners. The TCS naming scheme for HIP11704 and V583 follows the published schema (40,41).

### Coevolutionary model of TCS partners

Direct coupling analysis (13) is used in the coevolutionary analysis for the HK-RR dataset, which contains the collection of sequences from presumed coevolving TCS partners. In the DCA method, a global joint probability distribution was estimated to model the distribution of amino acids in each position along the HK-RR alignment.

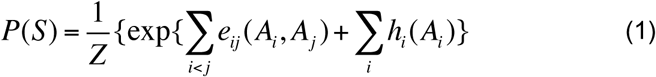

This Boltzmann distribution equation models the probability of a HK-RR sequence *S*. Indices *i* and *j* represent the *i*th and *j*th residue positions within the HK-RR alignment sequence. *Z* is a normalization factor (partition function). The detailed description of DCA model and the inference of the parameters, *e*_*ij*_(*A*_*i*_, *A*_*j*_) and *h*_*i*_(*A*_*i*_) can be found in previous study (13). The coupling strength parameter, *e*_*ij*_(*Ai, A*_*j*_), with 0<i⩽64 and 64<j⩽176, estimates the coevolutionary correlation or covariance strength between residues in HisKA domain and Response_reg domain. Each residue pair, ith residue and jth residue, has a 21 × 21 *e*_*ij*_ matrix, which contains coupling strength values for any amino acid pairing combination (selected from 20 amino acids and 1 gap). The local field parameter, *h*_*i*_(*A*_*i*_), is a 1 × 21 matrix and represents the preference of an amino acid at residue *i*.

### Hamiltonian score for binding specificity HK-RR pairs in *E. faecalis*

Protein sequences for HIP11704, OG1RF and V583 were downloaded from NCBI and were searched against HMM profile of HisKA domain (Pfam: PF00512) and Response_reg domain (Pfam: PF00072) by using hmmscan. HisKA domains and Response_reg domains of proteins from each strain were obtained and aligned. An energy function, Hamiltonian score (H(S)), was introduced to describe the coevolutionary energy of a sequence. The Boltzmann distribution can be written with the format of H(S) as:

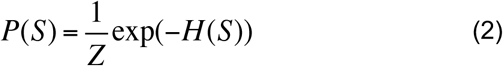

H(S) is:

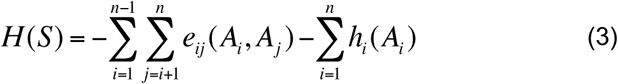

A specific energy function, binding specificity score Ĥ(S), was then calculated to determine the binding specificity between any HK and RR pair. Only the residual pairs at the binding interface were included in the calculation to focus on the binding specificity of HK and RR. We used a crystal structure (PDB ID: 3DGE) of HK and RR complex to provide the information of residues at the interface (distance ≤ 16 Å) of the HisKA and Response_reg functional domains. The Ĥ(S) of a given concatenated HK-RR sequence (S) is calculated as:

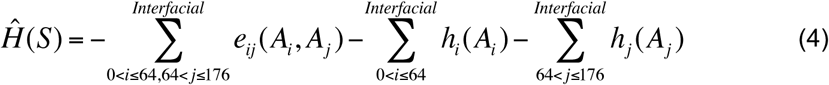

where *i* represents the residual position in HK and j represents the residual position in RR, where both are constrained by interfacial pairs. An Ĥ_0_(S) score representing the background noise for the same sequence was also calculated by using another set of *e*_*ij*_ and *h*_*i*_ parameters, which were estimated by using DCA with randomly paired and concatenated HK and RR sequences.

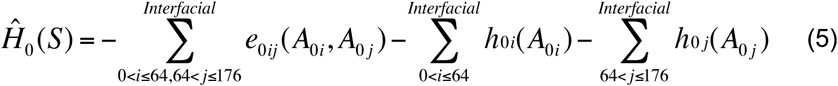

The difference between Ĥ(S) and Ĥ_0_(S), ΔĤ(S), was used to evaluate the binding specificity of any two HK and RR pairs. A more negative ΔĤ(S) score indicates stronger binding specificity.

### Antibiotic susceptibility test and growth curves for RNA-Seq

*E. faecalis* strains were grown in brain heart infusion (BHI) agar or broth overnight at 37°C. Antibiotic minimum inhibitory concentrations (MIC) were determined by broth microdilution. Antibiotics were serially diluted two-fold in BHI. 5 μL of overnight cultures of *E. faecalis* was diluted 1:1000 with 5 mL of fresh BHI before adding into wells. The plates were incubated in 37°C, recording the lowest antibiotic concentrations without growth as the MIC at 24 and 48 hours. The MICs obtained and used to design RNA-Seq experiments were 2048 μg/mL ceftriaxone for *E. faecalis* HIP11704 and 2048 μg/mL ceftriaxone for *E. faecalis* V583, 256 μg/mL vancomycin for V583, and 2048 μg/mL vancomycin for HIP11704 (Table 2).

**Table 2.** Median ceftriaxone and vancomycin MICs in BHI.

|  | Van Type | Ceftriaxone MIC<br>(µg/ml) | Vancomycin MIC<br>(µg/ml) |
| --- | --- | --- | --- |
| *E. faecalis* HIP11704 | vanA | 2048 | 2048 |
| *E. faecalis* V583 | vanB | 2048 | 256 |

For RNA-Seq experiments, overnight bacterial cultures were diluted to an OD_600nm_ of 0.01 with fresh BHI. The cultures were incubated at 37°C without agitation until the OD_600nm_ reached 0.5-0.6. 20 mL of the culture was added to an equal amount of BHI with vancomycin (256 μg/ml for V583 and 2048 μg/ml for HIP11704), ceftriaxone (2048 μg/ml), or BHI with no antibiotic supplementation (control). The growth curves were monitored by OD_600_ for 7 hours. For samples treated with ceftriaxone, bacterial samples were collected for RNA-Seq at 30 and 60 minutes post-treatment. With vancomycin, samples were collected at 15 and 45 minutes post-treatment. The difference in collection times seeks to adjust for differences in growth curves between antibiotic treatments (Figure S9). Three independent replicates were performed for each condition.

### RNA sequencing

RNA isolation, DNase treatment, and RNA cleanup were performed as previously described (24). RNA-Seq was performed at the University of Texas at Dallas Genome Center. RiboMinus Transcriptome Isolation kit and RiboMinus Concentration Module were used for rRNA depletion (ThermoFisher). Library preparation was then completed using Illumina Truseq Stranded mRNA library prep kit, where the mRNA selection step was skipped to yield total bacterial RNA for each sample. Then, total library sequencing was performed using single-read sequencing with a read length of 75 nucleotides using the Illumina NextSeq500.

### Transcriptomics analysis

RNA-sequencing data analysis was performed with CLC Genomics Workbench and DESeq2 R package (25). V583 and HIP11704 references were downloaded from NCBI (accession number: V583: AE016830.1, HIP11704: NZ_GG692622-59). Reads were aligned and counted to these references using default parameters after removing all the reads mapped to rRNAs and tRNAs in CLC Genomics Workbench. The unique read counts for each gene were normalized and then the fold change of expression level in treatment group versus control group was analyzed by R package, DESeq2. Differentially expressed genes were selected with the criteria of absolute value of log2 fold change larger than 1 and false discovery rate (FDR) adjusted p value <0.05.

### KEGG orthology (KO) analysis

The KO dataset for *E. faecalis* V583 was obtained from KEGG database (26,27). The differentially expressed genes (DEGs) of V583 were annotated with the genes’ functional orthologs by using the KO dataset. DEGs in HIP11704 were firstly mapped to the orthologous genes in V583 and then were searched against the KO dataset to be annotated.

### UMAP clustering analysis

Read counts for DEGs were imported into *R* statistical framework (44). Low read counts (<10) were filtered out. Counts for each test group were normalized and input to the uniform manifold approximation (UMAP) algorithm. UMAP clustering was performed using the cosine method with a minimum distance of 0.001 and nearest neighbors of 10. These parameters optimize clustering to give a balance of global and local information. Dbscan was used the identify different clusters with ε=0.6 for HIP11704 treated with ceftriaxone, ε=0.45 for HIP11704 treated with vancomycin, and ε=0.7 for V583 treated with ceftriaxone and vancomycin. The parameter ε was chosen to optimize clustering such that known interacting partners were placed in the same cluster where possible.

### Accession numbers

Illumina reads from RNA-Seq experiments have been deposited in the Sequence Read Archive under BioProject **PRJNA781303**.

## Acknowledgements

Funding from the University of Texas at Dallas (XJ, FM); National Institute of Health R35GM133631 (XJ, FM); National Science Foundation MCB-1943442 (CZ, FM); Cecil H. and Ida Green Chair in Systems Biology Science at the University of Texas at Dallas (KP).

## References

1. Bem AE, Velikova N, Pellicer MT, Baarlen P, Marina A, Wells JM. Bacterial histidine kinases as novel antibacterial drug targets. ACS Chem Biol. 2015 Jan 16;10(1):213–224.

2. Mitrophanov AY, Groisman EA. Signal integration in bacterial two-component regulatory systems. Genes Dev. 2008 Oct 1;22(19):2601–11.

3. Gao R, Mack TR, Stock AM. Bacterial response regulators: versatile regulatory strategies from common domains. Trends Biochem Sci. 2007 May;32(5):225–234.

4. Palmer KL, Gilmore MS. Multidrug-resistant enterococci lack CRISPR-cas. mBio. 2010 Oct 12;1(4):e00227–10.

5. Hong HJ, Hutchings MI, Buttner MJ; Biotechnology and Biological Sciences Research Council, UK. Vancomycin resistance VanS/VanR two-component systems. Adv Exp Med Biol. 2008;631:200–13.

6. Arthur M, Quintiliani R Jr. Regulation of VanA- and VanB-type glycopeptide resistance in enterococci. Antimicrob Agents Chemother. 2001 Feb;45(2):375–81.

7. Boyd JS, Cheng RR, Paddock ML, Sancar C, Morcos F, Golden SS. A Combined Computational and Genetic Approach Uncovers Network Interactions of the Cyanobacterial Circadian Clock. J Bacteriol. 2016 Aug 25;198(18):2439–47.

8. Skerker JM, Perchuk BS, Siryaporn A, Lubin EA, Ashenberg O, Goulian M, Laub MT. Rewiring the specificity of two-component signal transduction systems. Cell. 2008 Jun 13;133(6):1043–54.

9. Podgornaia AI, Casino P, Marina A, Laub MT. Structural basis of a rationally rewired protein-protein interface critical to bacterial signaling. Structure. 2013 Sep 3;21(9):1636–47.

10. Kellogg SL, Kristich CJ. Functional Dissection of the CroRS Two-Component System Required for Resistance to Cell Wall Stressors in Enterococcus faecalis. J Bacteriol. 2016 Mar 31;198(8):1326–36.

11. Koretke KK, Lupas AN, Warren PV, Rosenberg M, Brown JR. Evolution of two-component signal transduction. Mol Biol Evol. 2000 Dec;17(12):1956–70.

12. Cheng RR, Morcos F, Levine H, Onuchic JN. Toward rationally redesigning bacterial two-component signaling systems using coevolutionary information. Proc Natl Acad Sci U S A. 2014 Feb 4;111(5):E563–71.

13. Morcos F, Pagnani A, Lunt B, Bertolino A, Marks DS, Sander C, Zecchina R, Onuchic JN, Hwa T, Weigt M. Direct-coupling analysis of residue coevolution captures native contacts across many protein families. Proc Natl Acad Sci U S A. 2011 Dec 6;108(49):E1293–301.

14. Jana B, Morcos F, Onuchic JN. From structure to function: the convergence of structure based models and co-evolutionary information. Phys Chem Chem Phys. 2014 Mar 07; 16:6496–6507.

15. Morcos F, Jana B, Hwa T, Onuchic JN. Coevolutionary signals across protein lineages help capture multiple protein conformations. Proc Natl Acad Sci U S A. 2013 Dec 17;110(51):20533–8.

16. dos Santos RN, Morcos F, Jana B, Andricopulo AD, Onuchic JN. Dimeric interactions and complex formation using direct coevolutionary couplings. Sci Rep. 2015 Sep 04;5:13652.

17. Jiang XL, Martinez-Ledesma E, Morcos F. Revealing protein networks and gene-drug connectivity in cancer from direct information. Sci Rep. 2017 Jun 16;7(1):3739.

18. Zhou Q, Kunder N, De la Paz JA, Lasley AE, Bhat VD, Morcos F, Campbell ZT. Global pairwise RNA interaction landscapes reveal core features of protein recognition. Nat Commun. 2018 Jun 28;9(1):2511.

19. Dimas RP, Jiang XL, Alberto de la Paz J, Morcos F, Chan CTY. Engineering repressors with coevolutionary cues facilitates toggle switches with a master reset. Nucleic Acids Res. 2019 Jun 04;47(10):5449–5463.

20. El-Gebali S, Mistry J, Bateman A, Eddy SR, Luciani A, Potter SC, Qureshi M, Richardson LJ, Salazar GA, Smart A, Sonnhammer ELL, Hirsh L, Paladin L, Piovesan D, Tosatto SCE, Finn RD. The Pfam protein families database in 2019. Nucleic Acids Res. 2019 Jan 8;47(D1):D427–D432.

21. UniProt Consortium. UniProt: a worldwide hub of protein knowledge. Nucleic Acids Res. 2019 Jan 8;47(D1):D506–D515.

22. Potter SC, Luciani A, Eddy SR, Park Y, Lopez R, Finn RD. HMMER web server: 2018 update. Nucleic Acids Res. 2018 Jul 2;46(W1):W200–W204.

23. Hullahalli K, Rodrigues M, Palmer KL. Exploiting CRISPR-Cas to manipulate Enterococcus faecalis populations. Elife. 2017 Jun 23;6:e26664.

24. Hullahalli K, Rodrigues M, Nguyen UT, Palmer K. An Attenuated CRISPR-Cas System in Enterococcus faecalis Permits DNA Acquisition. mBio. 2018 May 1;9(3):e00414–18.

25. Love MI, Huber W, Anders S. Moderated estimation of fold change and dispersion for RNA-seq data with DESeq2. Genome Biol. 2014 Dec 05;15:550.

26. Kanehisa M, Goto S. KEGG: kyoto encyclopedia of genes and genomes. Nucleic Acids Res. 2000 Jan 1;28(1):27–30.

27. Kanehisa M, Sato Y, Furumichi M, Morishima K, Tanabe M. New approach for understanding genome variations in KEGG. Nucleic Acids Res. 2019 Jan 8;47(D1):D590–D595.

28. Howell A, Dubrac S, Noone D, Varughese KI, Devine K. Interactions between the YycFG and PhoPR two-component systems in Bacillus subtilis: the PhoR kinase phosphorylates the non-cognate YycF response regulator upon phosphate limitation. Mol Microbiol. 2006 Feb;59(4):1199–215.

29. Nakayama J, Cao Y, Horii T, Sakuda S, Akkermans AD, de Vos WM, Nagasawa H. Gelatinase biosynthesis-activating pheromone: a peptide lactone that mediates a quorum sensing in Enterococcus faecalis. Mol Microbiol. 2001 Jul;41(1):145–54.

30. Hancock LE, Perego M. The Enterococcus faecalis fsr two-component system controls biofilm development through production of gelatinase. J Bacteriol. 2004 Sep;186(17):5629–39.

31. Muller C, Le Breton Y, Morin T, Benachour A, Auffray Y, Rincé A. The response regulator CroR modulates expression of the secreted stress-induced SalB protein in Enterococcus faecalis. J Bacteriol. 2006 Apr;188(7):2636–45.

32. Le Breton Y, Muller C, Auffray Y, Rincé A. New insights into the Enterococcus faecalis CroRS two-component system obtained using a differential-display random arbitrarily primed PCR approach. Appl Environ Microbiol. 2007 Jun;73(11):3738–41.

33. Kristich CJ, Djorić D, Little JL. Genetic basis for vancomycin-enhanced cephalosporin susceptibility in vancomycin-resistant enterococci revealed using counterselection with dominant-negative thymidylate synthase. Antimicrob Agents Chemother. 2014;58(3):1556–64.

34. Kristich CJ, Rice LB, Arias CA. Enterococcal Infection—Treatment and Antibiotic Resistance. 2014 Feb 6. In: Gilmore MS, Clewell DB, Ike Y, et al., editors. Enterococci: From Commensals to Leading Causes of Drug Resistant Infection [Internet]. Boston: Massachusetts Eye and Ear Infirmary; 2014-

35. Ahmed MO, Baptiste KE. Vancomycin-Resistant Enterococci: A Review of Antimicrobial Resistance Mechanisms and Perspectives of Human and Animal Health. Microb Drug Resist. 2018 Jun;24(5):590–606.

36. Palmer KL, Carniol K, Manson JM, Heiman D, Shea T, Young S, Zeng Q, Gevers D, Feldgarden M, Birren B, Gilmore MS. High-quality draft genome sequences of 28 Enterococcus sp. isolates. J Bacteriol. 2010 May;192(9):2469–70.

37. Sahm DF, Kissinger J, Gilmore MS, Murray PR, Mulder R, Solliday J, Clarke B. In vitro susceptibility studies of vancomycin-resistant Enterococcus faecalis. Antimicrob Agents Chemother. 1989 Sep;33(9):1588–91.

38. Paulsen IT, Banerjei L, Myers GS, Nelson KE, Seshadri R, Read TD, et al. Role of mobile DNA in the evolution of vancomycin-resistant Enterococcus faecalis. Science. 2003 Mar 28;299(5615):2071–4.

39. Gold OG, Jordan HV, van Houte J. The prevalence of enterococci in the human mouth and their pathogenicity in animal models. Arch Oral Biol. 1975 Jul;20(7):473–7.

40. Hancock LE, Perego M. Systematic inactivation and phenotypic characterization of two-component signal transduction systems of Enterococcus faecalis V583. J Bacteriol. 2004 Dec;186(23):7951–8.

41. Hancock L, Perego M. Two-Component Signal Transduction in Enterococcus faecalis. J Bacteriol. 2002 Nov 01;184:5819–5825.

42. Tiwari S, Jamal SB, Hassan SS, Carvalho PVSD, Almeida S, Barh D, Ghosh P, Silva A, Castro TLP, Azevedo V. Two-Component Signal Transduction Systems of Pathogenic Bacteria As Targets for Antimicrobial Therapy: An Overview. Front Microbiol. 2017 Oct 10;8:1878.

43. Sinner C, Ziegler C, Jung YH, Jiang X, Morcos F. ELIHKSIR Web Server: Evolutionary Links Inferred for Histidine Kinase Sensors Interacting with Response Regulators. Entropy (Basel). 2021 Jan 30;23(2):170.

44. R Core Team. R: A Language and Environment for Statistical Computing. R Foundation for Statistical Computing (Vienna, Austria) 2020:https://www.R-project.org/

